# Expanding the marine range of the endangered black-capped petrel *Pterodroma hasitata*: Occurrence in the northern Gulf of Mexico and conservation implications

**DOI:** 10.1101/2021.01.19.427288

**Authors:** Patrick G.R. Jodice, Pamela E. Michael, Jeffrey S. Gleason, J. Christopher Haney, Yvan G. Satgé

**Affiliations:** U.S. Geological Survey, South Carolina Cooperative Fish & Wildlife Research Unit, Clemson University, Clemson, South Carolina 29634, USA; South Carolina Cooperative Fish & Wildlife Research Unit, Clemson University, Clemson, South Carolina 29634, USA; U.S. Fish and Wildlife Service, Gulf Restoration Team, Chiefland, Florida 32626, USA; Terra Mar Applied Sciences, LLC, Washington, D.C. 20012, USA

**Keywords:** Black-capped petrel, *Pterodroma hasitata*, Vessel surveys, Gulf of Mexico, Range

## Abstract

The black-capped petrel (*Pterodroma hasitata*) is an endangered seabird endemic to the western north Atlantic. Although estimated at ~ 1,000 breeding pairs, only ~ 100 nests have been located at two sites in Haiti and three sites in the Dominican Republic. At sea, the species primarily occupies waters of the western Gulf Stream in the Atlantic and the Caribbean Sea. Due to limited data, there is currently not a consensus on the marine range of the species. There are several maps in use for the marine range of the species and these differ with respect to the north, south, and eastward extent of the range. None of these maps, however, includes the Gulf of Mexico. Here, we report on observations of black-capped petrels during two vessel-based survey efforts throughout the northern Gulf of Mexico from July 2010 - July 2011, and from April 2017 - September 2019. During the 558 days and 54.7 km of surveys from both efforts we tallied 40 black-capped petrels. Most observations occurred in the eastern Gulf, although birds were observed over much of the east-west and north-south footprint of the survey area. Predictive models indicated that habitat suitability for black-capped petrels was highest in areas associated with dynamic waters of the Loop Current, similar to habitat used along the western edge of the Gulf Stream in the western north Atlantic. We suggest that the range for black-capped petrels be modified to include the entire northern Gulf of Mexico although distribution may be more clumped in the eastern Gulf and patchier elsewhere. It remains unclear, however, which nesting areas are linked to the Gulf of Mexico.

## INTRODUCTION

One of the most basic attributes needed for conservation planning for wildlife at the species level is a current map of annual range (Noss et al. 1997). Without such information, conservation threats cannot be understood and our ability to prioritize mitigation measures, conservation actions, and research is limited (Underhill & Gibbons 2002, Gibbons et al. 2010, Liminana et al. 2015). For example, Cooper et al. (2019) recently revised the range for the endangered Kirtland’s warbler (*Setophaga kirlandii*), and as a result of a redefined range the authors identified additional threats to, and research priorities for, that species. For many threatened and endangered avian species, the primary tools used to assess or refine the range are location-specific surveys and individual-based tracking. Surveys for endangered species typically use point-counts or transects to map occurrence of a species within a focal area. These focal areas may be chosen because they harbor some conservation threat or because the area has yet to be surveyed, but appears suitable for occupancy (Cooper et al. 2019, Ortega-Alvarez et al. 2020). Surveys can also be conducted using newer technologies such as audio recording units or camera trapping, both of which can enhance sampling effort particularly in remote locations (Beirne et al. 2017, Schroeder & McRae 2020). The range of a species can also be refined based on data obtained from individual tracking efforts, and such data also can be used to locate previously unidentified breeding or nonbreeding areas (Kanai et al. 2002, McCloskey et al. 2018). Advantages of the latter approach are that areas not previously considered or too remote to survey may be ‘discovered’, residency time within an area may be determined, and interactions of individuals with conservation threats may be identified (Jodice et al. 2015, Lamb et al. 2018, Phillips et al. 2018). More recently, citizen-science data, such as eBird (www.ebird.org), have been used to add detail to the range of a species by providing unique or rare sightings (Cooper et al. 2019).

Defining the range of pelagic seabirds presents unique challenges given that the annual cycle is dominated by time spent in remote locations at sea. During the breeding season the spatial extent of the foraging range of pelagic seabirds can extend 100s to 1000s of kilometers, while during the nonbreeding season individuals can range over entire ocean basins (Jodice & Suryan 2010, Phillips et al. 2007, Rayner et al. 2010). Their extensive ranging behavior exposes individuals to a wide array of marine threats including oil and gas activity (Haney et al. 2017), bycatch mortality from fisheries operations (Anderson et al. 2011), and pollution events (Provencher et al. 2020). These threats often occur across multiple political boundaries or in international waters, the latter of which can be poorly monitored and regulated (Jodice & Suryan 2010). Therefore, range maps for pelagic seabirds are often lacking in detail, despite this group having a high proportion of threatened and endangered species (Croxall et al. 2012, Dias et al. 2019). Because range maps for seabirds often lack resolution, spatially explicit assessments of conservation threats also can be deficient for this group of birds (Oppel et al. 2012, Jodice et al. 2019). The combination of poorly defined marine ranges and transboundary marine threats adds to the conservation challenges faced by many species of pelagic seabirds.

Among seabirds, one of the least studied and most threatened groups are the gadfly petrels (*Pterodroma* spp.), which often nest on remote islands and inhabit pelagic waters both near and distant to their nesting areas. The Atlantic Ocean supports 11 species of gadlfy petrel (Ramos et al. 2017), two of which are extant in the western North Atlantic (*P. cahow* and *P. hasitata*). Here, we focus on black-capped petrel (*Pterodroma hasitata*; also known locally as Diablotín). Simons et al. (2013) provide a thorough review of the biology and conservation of the species and Satgé et al. (2020) of nesting habitat relationships. This species is considered globally endangered (BirdLife International 2020; hereafter, any reference to the species as endangered refers to its global status) and is under consideration for listing as Threatened with 4(d) under the U.S. Endangered Species Act (USFWS 2018; 83 FR 50560). Black-capped petrels nest in the understory of montane forests at 1,500 – 2,000 m above sea level in burrows and crevices. The breeding season occurs primarily from February – July with birds dispersing at sea thereafter (Simons et al. 2013). The black-capped petrel was considered extinct in the mid-1900s but was rediscovered in 1963 when nests were located in the Massif de la Selle of southeastern Haiti (Wingate 1964). Since that time, ~100 nests have been found (n = two sites in Haiti, n = three sites in the Dominican Republic; Fig. 1). The nesting area in Valle Nuevo in the Cordillera Central of the Dominican Republic was documented to support nesting only as recently as 2018. Nesting is suspected in Dominica and Cuba but has yet to be confirmed.

**Fig. 1.**
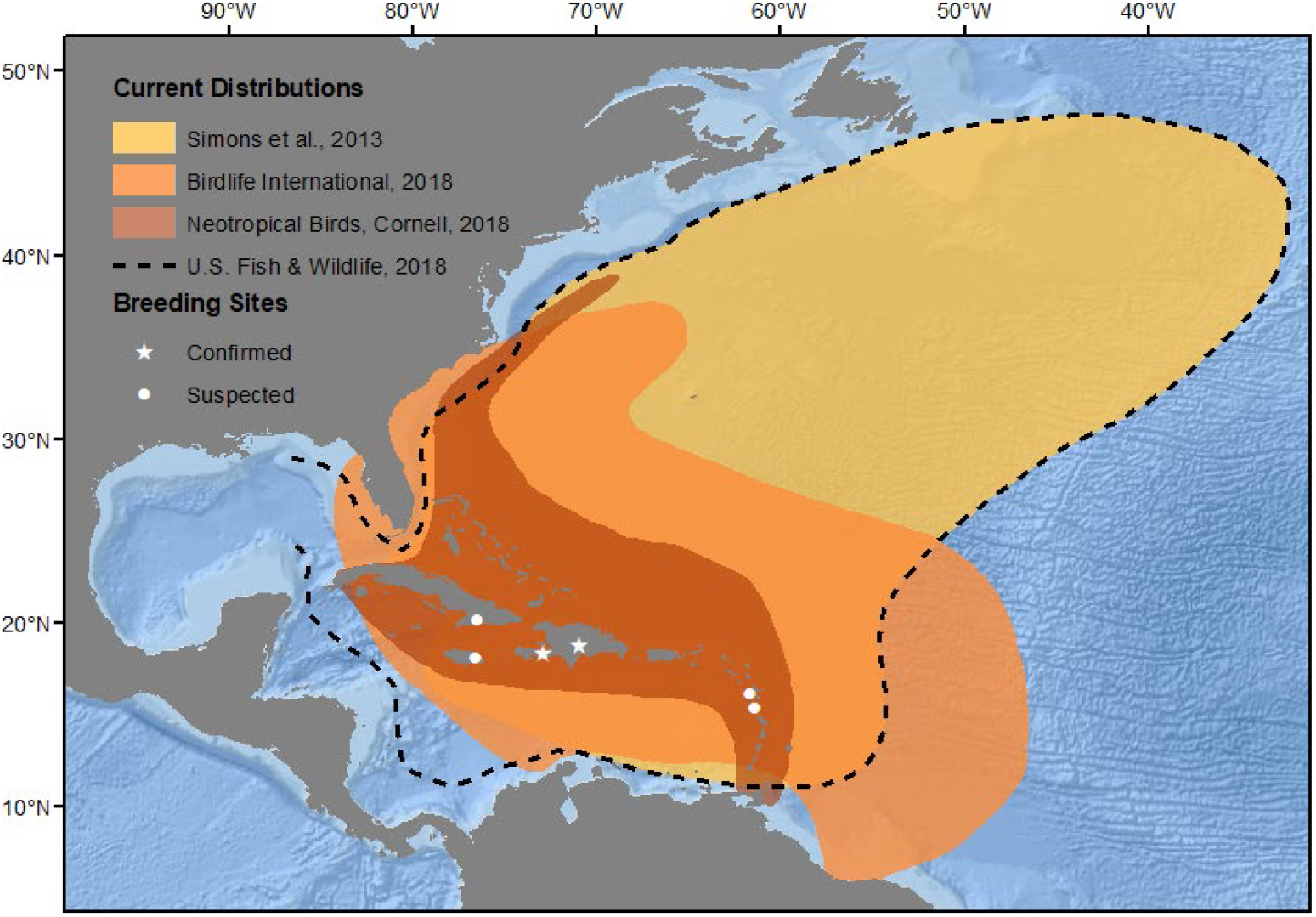
Breeding locations (known and suspected) and marine range of black-capped petrel (*Pterodroma hasitata*). Breeding sites are labeled as suspected (e.g., evidence of black-capped petrel presence based on audio or radar surveys) and documented. The marine range differs among four primary sources and each is displayed for reference. Credit for base map: ESRI, Garmin, GEBCO, NOAA, NGDC, and other contributors.

At-sea, most of what is known about the range of the species is based on observations from vessel-based surveys in the western north Atlantic (Haney 1987, Simons et al. 2013, Winship et al. 2018) and recent efforts to track individuals (Jodice et al. 2015, Satgé et al. 2019). These data sets primarily place the range of the species in the western north Atlantic between ~ 30 - 40 degrees latitude and west of the Gulf Stream, although waters east of the Gulf Stream and in the Caribbean Sea also were highlighted as use areas via tracking data. Based on these data, several sources have developed range maps for the species and while each differs slightly, all focus on waters west of the Gulf Stream and none include definitive use of the Gulf of Mexico (Fig. 1). The species also occurs in both a light and dark color morph (Howell & Patteson 2008); it is unclear, however, if the ranges of these two morphs are similar or disparate either spatially or temporally.

Herein, we provide new documentation for expanding the marine range of the endangered black-capped petrel to include the northern Gulf of Mexico. We used data from two vessel-based survey efforts that occurred throughout the northern Gulf and used a machine-learning modeling approach to describe basic habitat relationships in this basin. These data represent a substantial refinement of the marine range of a globally endangered species and do so in a region with extensive offshore oil and gas exploration and development.

## METHODS

### At-sea surveys

Observations of black-capped petrels in the Gulf of Mexico (hereafter, GoM or Gulf) were collected during vessel-based surveys for pelagic seabirds conducted as part of two survey programs (Table S1). Surveys to support the post-spill injury assessment for the *Deepwater Horizon* Oil Spill Natural Resources Damage Assessment (NRDA) were designed to record occurrences of seabirds and assess mortality and visible oiling (hereafter NRDA cruises; Haney 2011). NRDA cruises (*n* = 27; see Haney et al. 2019 for a detailed description) were conducted in the northern GoM within the U.S. Exclusive Economic Zone (EEZ) from July 2010 - July 2011 by experienced seabird observers (Fig. 2a). Surveys were conducted across 283 days and ~15,300 km of transects (Table S1). Surveys to support the Gulf of Mexico Marine Assessment Program for Protected Species (hereafter GoMMAPPS cruises) sought to model the distribution of seabirds, marine mammals, and sea turtles in the northern Gulf in relation to oil and gas activities among planning areas delineated by the U.S. Bureau of Ocean Energy Management (BOEM; https://www.boem.gov/regions/gulf-mexico-ocs-region). GoMMAPPS surveys (*n* = 20) were conducted in the northern GoM within the U.S. EEZ from April 2017 - September 2019 by experienced seabird observers (Fig. 2b). GoMMAPPS surveys were conducted from National Oceanic and Atmospheric Administration (NOAA) vessels of opportunity that were designed to survey for marine mammals or to collect fisheries/plankton data. Surveys were conducted during 275 days and ~39,400 km of transects (Table S1).

**Fig. 2.**
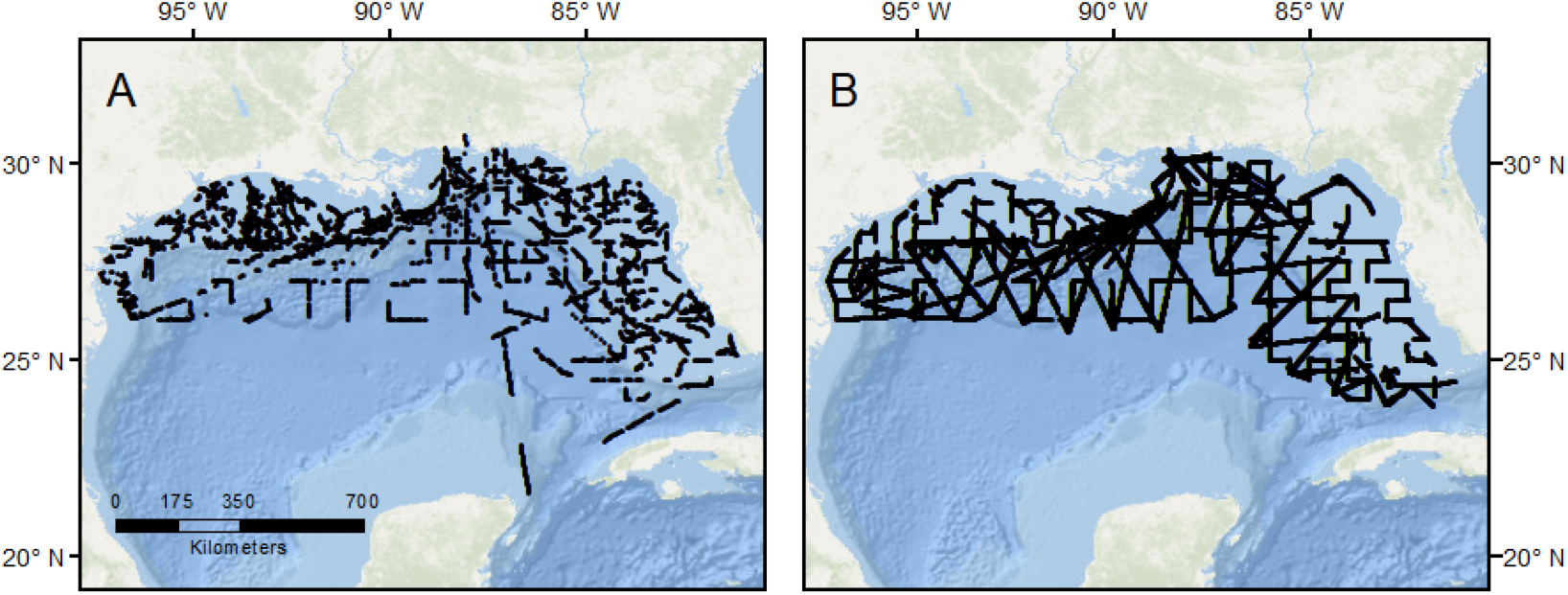
Spatial footprint of research cruises from two research programs in the northern Gulf of Mexico from which black-capped petrels (*Pterodroma hasitata*) were observed. (A) Surveys conducted in 2010 and 2011 were a component of the post-spill *Deepwater Horizon* Natural Resources Damage Assessment (NRDA). (B) Surveys conducted in 2017 – 2019 were a component of the Gulf of Mexico Marine Assessment Program for Protected Species (GoMMAPPS). Credit for base map: ESRI, Garmin, GEBCO, NOAA, NGDC, and other contributors.

Data collection during each survey program followed a standardized protocol for collection of marine fauna at sea (e.g., Tasker 2004). Briefly, trained observers surveyed for seabirds from a platform onboard the vessel situated ~13-15 m above the sea surface. While the vessel was underway at a speed ≥ ~11 km/h, the observer used ≥ 10x binoculars to identify to the lowest taxonomic level all sitting and flying seabirds within view. Observations were made from the side of the ship with the least glare (i.e., focal side). Following standard protocols, we recorded all seabirds within a 90° forward-facing arc. Relatively low densities and good observation conditions in the GoM, however, generally allowed species-specific identification and accurate counts beyond the typical 300 m width, out to ~ 700 m from the ship. Because black-capped petrels are surface foragers we did not need to account for time below the surface or observations missed during diving. During NRDA cruises, observations of seabirds were recorded manually (i.e., paper, voice recording) along with the exact time and later synchronized with the position of the ship as recorded by GPS at 10-minute intervals. During GoMMAPPS cruises, observations were recorded in real-time using the software package SEEBIRD (Ballance & Force 2016). For each entry, the observer recorded the species, number of individuals, distance bin or approximate distance, associations with other species, behavior, flight direction, flight height, flight angle, and when possible age, sex, and plumage. SEEBIRD records date, time, and GPS location at the time the record is initially ‘opened’ via direct connection with the ships navigation system.

To complement our data, we also sought other records of black-capped petrels in the Gulf of Mexico. We reviewed published literature and reports from seabird surveys conducted during the GulfCet I and II programs (Davis and Fargion 1995, Ribic et al. 1997, Davis et al. 2000). We also reviewed compilations that included seabirds (Duncan and Havard 1980, Clapp et al. 1982) and a monograph focused on the black-capped petrel (Simons et al. 2013). Lastly, we searched eBird for records of black-capped petrels.

### Modelling approach

We modeled the probability of occurrence of black-capped petrels based on habitat suitability in the northern Gulf of Mexico using the maximum entropy approach in Program Maxent (Version 3.4.2; https://biodiversityinformatics.amnh.org/open_source/maxent/; Phillips et al. 2006). Briefly, Maxent is a machine learning technique that estimates the probability of occurrence of a species across a specified area based on observations and a set of covariates (i.e., predictor variables that represent habitat conditions). Maxent performs well at relatively low sample sizes (i.e., *n* < 100 observations) by utilizing a presence-background algorithm that is less sensitive to sample size compared to other approaches used to model species distributions (Phillips et al. 2006, Wisz et al. 2008). We chose to model only presence records (i.e., as opposed to modeling both presence and pseudo-absence data) due to dissimilarities in the documentation of survey effort between NRDA and GoMMAPPS which prevented a fully standardized and comparable description of observation effort.

Using the Maxent interface, we estimated the probability of occurrence of black-capped petrels (based on habitat suitability, from 0 to 1) within 10,000 random background pixels across the entire Gulf of Mexico (i.e., “cloglog” output in Maxent). Data were modelled at a spatial resolution of 4.67 km based on the finest resolution available across the selected environmental data (see below for details). As observations occurred in only a portion of the study area, we applied “clamping” which, in Maxent, assumes that covariates from background pixels with values outside of the range of those from the training data can occur, but at low probabilities (i.e., at the tail end of the distribution; Philips et al. 2006). Clamping thus reduces the potential for predicting a high probability of occurrence in areas with covariate values well outside of those in the training data. We assessed model performance by separating the observations into randomly selected training and testing datasets (75%/25% split, respectively). We applied the model to the test data and used the area under the receiver operating characteristics curve (AUC) to quantify the predictive power of the model, where an AUC of 0.5 indicates no predictive power and an AUC of 1 indicates perfect discrimination (Bradley 1997). To balance the probability of incorrectly identifying suitable and unsuitable areas, we used the equal test sensitivity and specificity threshold (Cantor et al. 1999, Liu et al. 2005). We characterized the permutation importance of each covariate (the sensitivity of the model to a given covariate, holding all other covariates constant) using a jackknife procedure, a resampling approach to estimate variance and bias (Efron 1992).

We modeled observations of black-capped petrels in relation to static and dynamic environmental variables which were selected based on habitat relationships described for black-capped petrels in the Gulf Stream of the western North Atlantic (Haney 1987, Winship et al. 2018) and on other surface feeding seabirds in the Gulf of Mexico (Poli et al. 2017). Each dynamic variable was calculated as the temporal average based on daily dynamic variables for each date when a black-capped petrel was observed, weighted by the number of presence records of petrels on a given day. We obtained the daily dynamic variables of sea-surface temperature, sea-surface salinity, sea-surface height, and surface current velocity (eastward (*u*) and northward(*v*)) from the Hybrid Coordinate Ocean Model (HYCOM; Chassignet et al. 2009, Metzger et al. 2017). Surface current velocity was also used to calculate absolute current strength and current direction, each of which were subsequently included in the Maxent model as covariates. Although we considered including chlorophyll *a* as a predictor variable, spatial gaps in coverage would have resulted in the omission of observations of petrels. Preliminary assessments revealed that including chlorophyll *a* did not improve model performance and we therefore excluded chlorophyll *a* from subsequent analyses. Lastly, we included average depth as calculated from the SMRT30+ version 6.0 30 arc second dataset (Becker et al 2009). We aggregated each variable to the coarsest native spatial resolution available across all variables (~4.67 km). Therefore, the spatial resolution of the subsequent model (i.e., the resolution at which occurrence probability can be interpreted) is 4.67 km × 4.67 km which is comparable to similar data sets from vessel-based surveys in the western north Atlantic (e.g., Winship et al. 2018). The degree of covariance and correlation between environmental covariates was assessed using the Band Collection Statistics tool in ArcMap 10.8. None of the covariates exceeded the Pearson’s correlation coefficient of |0.75| threshold for exclusion. Thus, all covariates were retained. Unless otherwise noted, covariates were processed in R (R Development Core Team 2019).

To develop a proposed range within the Gulf of Mexico for black-capped petrels we used utilization distributions (UD) with kernel density estimation. We computed the 90% UD contour for all observation records from the NRDA and GoMMAPPS systematic surveys (package adehabitat in R; Calenge 2006). We then constructed a minimum convex polygon (MCP) that encompassed the areas represented by the utilization distribution contours and propose that area bounded by the MCP as the proposed range.

## RESULTS

### Previous records in the Gulf of Mexico

We searched published literature and reports for previous records of the species in the Gulf of Mexico (Table 1). Neither systematic surveys nor compilations of records noted any definitive observations of black-capped petrels at sea in the Gulf from ~ 1900 – 2010. Approximately 9-11 opportunistic records were reported in the mid-1990s and 2010s (e.g., birding records). Two of these birding records, in November 2016 and January 2017 and each occurring ~220 km southeast of Galveston Bay, Texas (~ 27.5°, −94.3°), are the only records of black-capped petrels in the Gulf during those months of the year (see below). Other efforts to summarize observations of black-capped petrels at-sea have a restricted range (e.g., Leopold et al. 2019 in the Caribbean Sea, Winship et al., 2018 in the Atlantic) and therefore do not include Gulf waters.

**Table 1.**
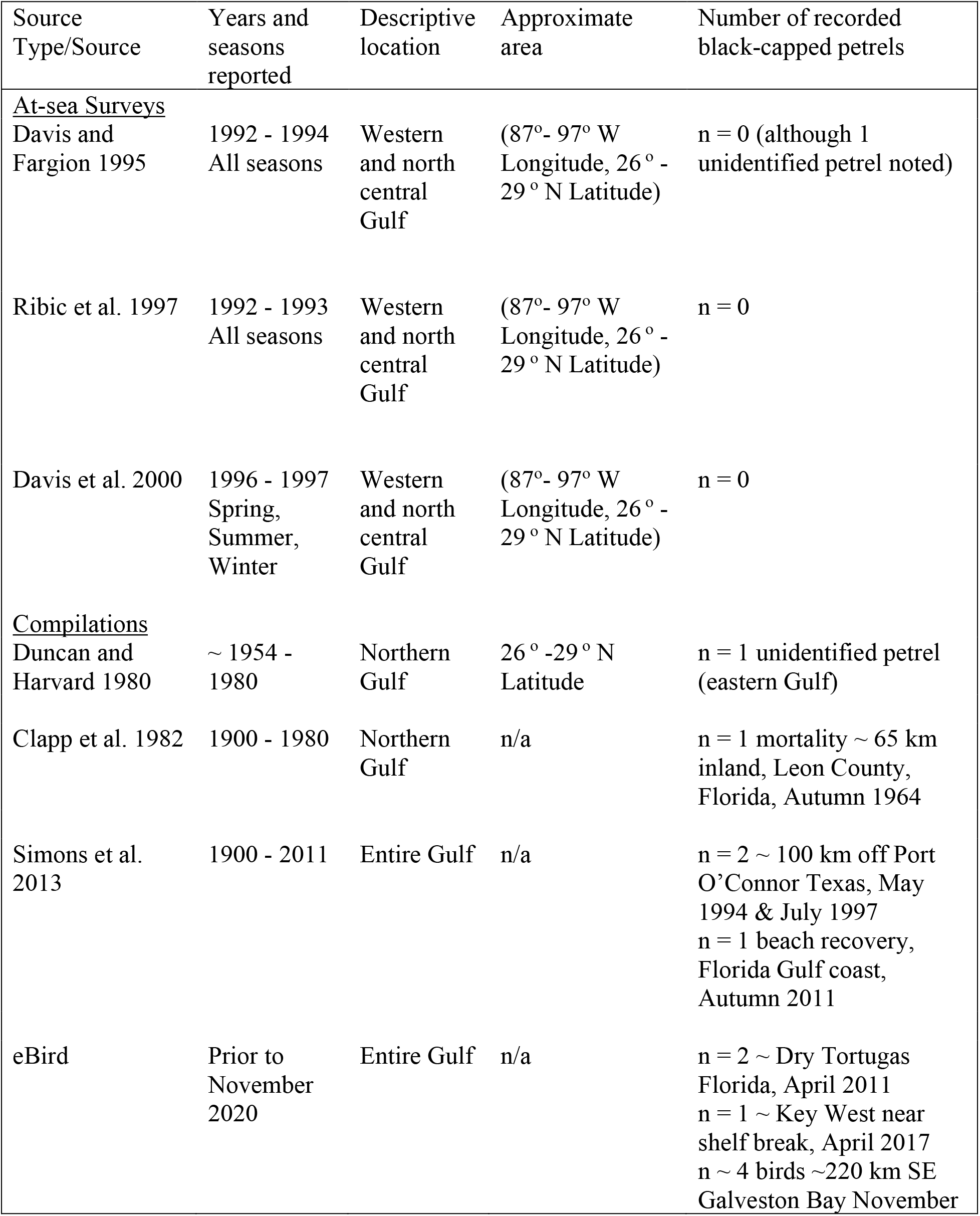
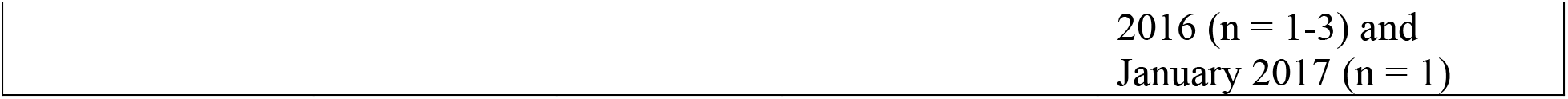
Summary of records of black-capped petrels (*Pterodroma hasitata*) for the Gulf of Mexico not including surveys from the post-spill Deepwater Horizon Natural Resources Damage Assessment (NRDA) or the Gulf of Mexico Marine Assessment Program for Protected Species (GoMMAPPS).

### Survey data

Observations made on NRDA cruises totaled nine black-capped petrels (Table 2, Fig. 3); six as singletons and one observation of three birds. One light-morph individual was observed in July 2010 in the eastern Gulf. Eight of the petrels observed during NRDA cruises occurred in waters east of the Mississippi River delta, while one bird was observed just west of the delta. Seven of the eight observations occurred in waters over the continental shelf break and slope. The southern extent of observations occurred slightly south and west of the western extent of the Florida Keys. We observed petrels in February - May, and July - September. Petrels were not observed during cruises in October – December, or in June.

**Table 2.**
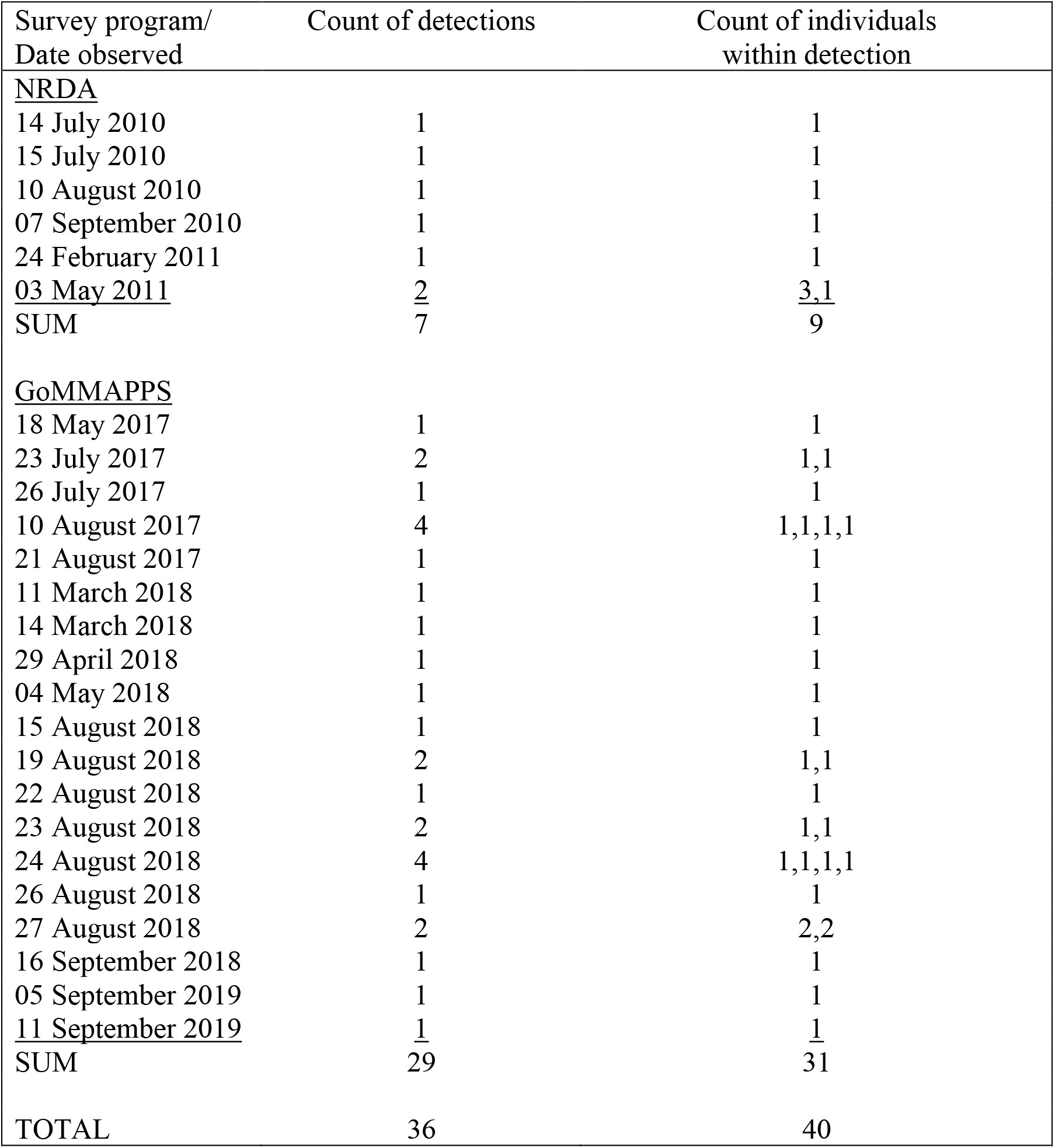
Black-capped petrels (*Pterodroma hasitata*) observed during research cruises from two research programs in the northern Gulf of Mexico. Surveys conducted in 2010 and 2011 were a component of the post-spill *Deepwater Horizon* Natural Resources Damage Assessment (NRDA). Surveys conducted in 2017 – 2019 were a component of the Gulf of Mexico Marine Assessment Program for Protected Species (GoMMAPPS). Detections refer to an observation of ≥ 1 black-capped petrel.

| Survey program/<br>Date observed | Count of detections | Count of individuals<br>within detection |
| --- | --- | --- |
| <u>NRDA</u> |  |  |
| 14 July 2010 | 1 | 1 |
| 15 July 2010 | 1 | 1 |
| 10 August 2010 | 1 | 1 |
| 07 September 2010 | 1 | 1 |
| 24 February 2011 | 1 | 1 |
| <u>03 May 2011</u> | <u>2</u> | <u>3,1</u> |
| SUM | 7 | 9 |
| <u>GoMMAPPS</u> |  |  |
| 18 May 2017 | 1 | 1 |
| 23 July 2017 | 2 | 1,1 |
| 26 July 2017 | 1 | 1 |
| 10 August 2017 | 4 | 1,1,1,1 |
| 21 August 2017 | 1 | 1 |
| 11 March 2018 | 1 | 1 |
| 14 March 2018 | 1 | 1 |
| 29 April 2018 | 1 | 1 |
| 04 May 2018 | 1 | 1 |
| 15 August 2018 | 1 | 1 |
| 19 August 2018 | 2 | 1,1 |
| 22 August 2018 | 1 | 1 |
| 23 August 2018 | 2 | 1,1 |
| 24 August 2018 | 4 | 1,1,1,1 |
| 26 August 2018 | 1 | 1 |
| 27 August 2018 | 2 | 2,2 |
| 16 September 2018 | 1 | 1 |
| 05 September 2019 | 1 | 1 |
| <u>11 September 2019</u> | <u>1</u> | <u>1</u> |
| SUM | 29 | 31 |
| TOTAL | 36 | 40 |

**Fig. 3.**
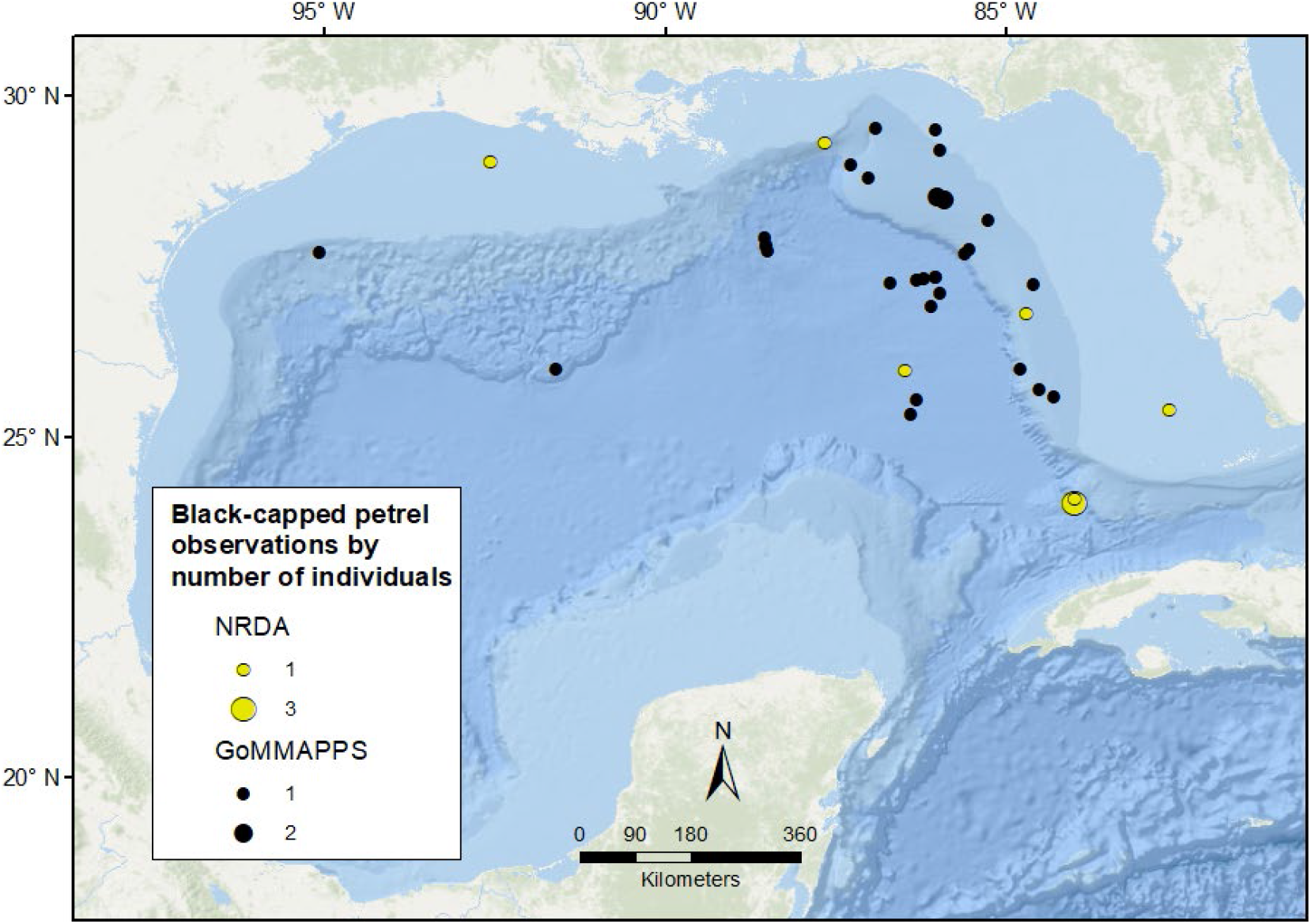
Locations of black-capped petrels (*Pterodroma hasitata*) observed during research cruises in the northern Gulf of Mexico. Surveys conducted in 2010 and 2011 were a component of the post-spill *Deepwater Horizon* Natural Resources Damage Assessment (NRDA). Surveys conducted in 2017 – 2019 were a component of the Gulf of Mexico Marine Assessment Program for Protected Species (GoMMAPPS). Credit for base map: ESRI, Garmin, GEBCO, NOAA, NGDC, and other contributors.

Observations made on GoMMAPPS cruises totaled 31 black-capped petrels, 27 of which included single birds and two observations of two birds (Table 2, Fig. 3). Three of the petrels observed on GoMMAPPS cruises were classified as light-morph individuals (March and August 2018 in the eastern Gulf). Most observations of black-capped petrels occurred in the eastern Gulf, east of −88° Longitude, although birds were observed over much of the east-west and north-south footprint of the survey area. We observed petrels in March - May and July - September. We did not observe birds on cruises in January - February, June, or October.

### Predictive Models

Our predictive model was relatively robust and generated an average AUC value of 0.909 for the training data set and 0.775 for the testing data set, compared to a maximum possible AUC of 0.877 (based on data being drawn from the Maxent distribution itself; Phillips et al. 2006). Maxent identified three general areas with relatively higher habitat suitability for black-capped petrels (Fig. 4). The most extensive of these occurs just west of the Florida shelf and extends along the north-south length of the Florida peninsula. Modeled habitat suitability also was higher south of the Mississippi River delta and within a narrow east-west band paralleling much of the Texas and Louisiana continental slope. Areas of lower habitat suitability include shelf/slope and pelagic waters in the central Gulf.

**Fig. 4.**
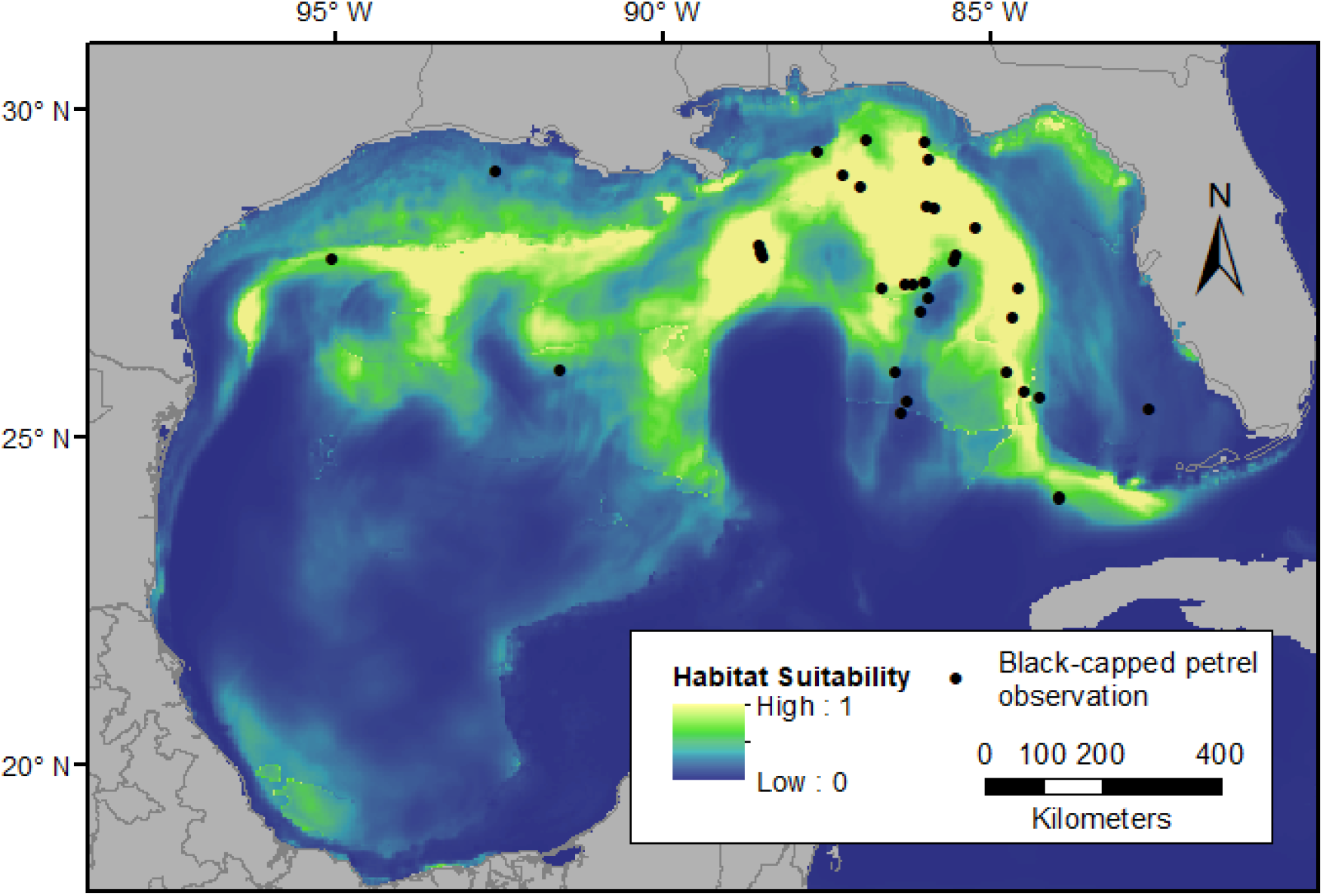
Predicted probability of black-capped petrel (*Pterodroma hasitata*) occurrence with observations from the post-spill Deepwater Horizon Natural Resources Damage Assessment (NRDA) and the Gulf of Mexico Marine Assessment Program for Protected Species (GoMMAPPS) surveys overlaid. Blue shades indicate a very small probability of occurrence while yellow shades indicate a high probability of occurrence based on habitat suitability. Credit for base map: ESRI, Garmin, GEBCO, NOAA, NGDC, and other contributors.

For black-capped petrels in the Gulf of Mexico, sea-surface salinity was the most important predictor of habitat suitability followed by the direction of the current and sea-surface height (Table 3). Habitat suitability was predicted to peak at moderate values of sea-surface height (Fig. 5a), to increase as currents were more eastward to southward (Fig. 5b), and to increase until a threshold with increasing salinity (Fig. 5c). Sea-surface salinity had the greatest permutation importance, with current direction and sea-surface height having less importance (Table 3).

**Table 3.** Relative contribution of environmental variables to habitat suitability of black-capped petrels (*Pterodroma hasitata*) in the northern Gulf of Mexico. See Methods for description of % contribution and permutation importance. Permutation importance sums to 100 across all variables.

| Variable | % contribution | Permutation importance |
| --- | --- | --- |
| Sea-surface salinity | 41.1 | 31.3 |
| Current direction | 24.2 | 0.4 |
| Sea-surface height | 18.5 | 10.5 |
| Bathymetry | 9.7 | 26.6 |
| Current velocity: v (northward) | 6.2 | 31.1 |
| Absolute current strength | 0.2 | 0.0 |
| Current velocity: u (eastward) | 0.1 | 0.0 |
| Sea-surface temperature | 0.0 | 0.0 |

**Fig. 5.**
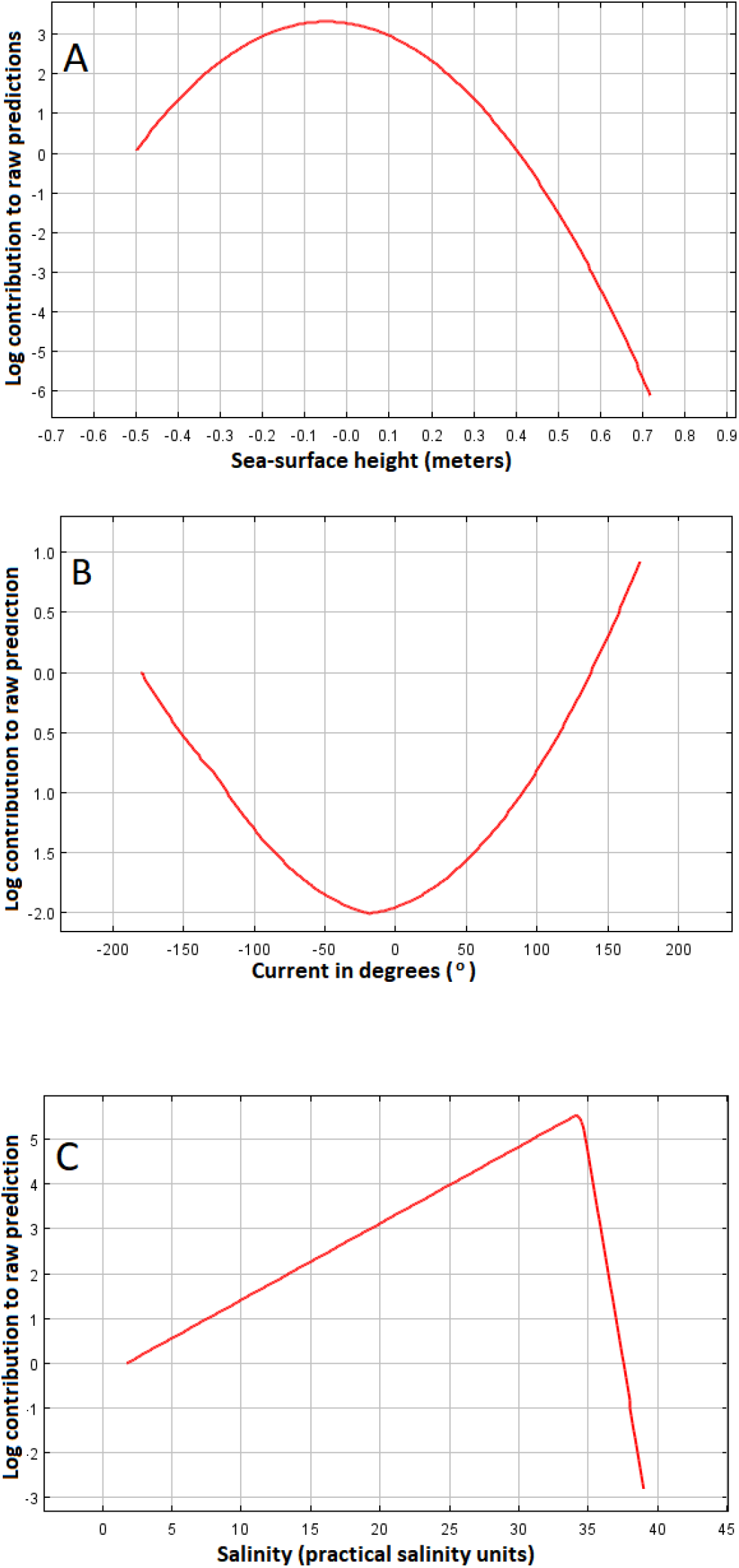
Response curves of habitat suitability of black-capped petrels (*Pterodroma hasitata*) in the northern Gulf of Mexico to (A) sea surface height, (B) current direction, and (C) sea surface salinity based on models developed in Maxent. Habitat suitability increases along the y axis (i.e., relatively higher probability of occurrence).

### DISCUSSION

Since the early 2000s, most effort invested in improving our understanding of the range of the black-capped petrel has been focused on locating nesting areas and detailing their areal extent. The marine range, in contrast, has been broadly accepted as being focused in waters of the Caribbean Sea (i.e., nearby known nesting areas) and along the western edge of the Gulf Stream in the western North Atlantic (Simons et al. 2013). This range is based on observations from vessel-based surveys focused primarily in the mid- and South-Atlantic bights of the U.S. that date back to the 1980s (Simons et al. 2013). Recent efforts to model the habitat preference of the species to support marine spatial planning in the western North Atlantic have also emphasized the western edge of the Gulf Stream and waters of the South Atlantic Bight as the primary marine range (Winship et al. 2018). Furthermore, tracking data from individuals tagged at nests in the Dominican Republic found that while waters of the southern Caribbean Sea and east of the Gulf Stream were used regularly during chick-rearing and post-breeding, waters of the Gulf of Mexico were not utilized (Jodice et al. 2015). Therefore, the Gulf of Mexico has yet to be recognized as a regular component of the marine range of the species.

Consequently, results from the NRDA and GoMMAPPS surveys have substantially revised our understanding of the marine range of the black-capped petrel. While prior data reviews and systematic vessel-based surveys in the Gulf failed to detect the species at-sea, we tallied 40 individuals, including light morph birds, across both efforts. Most observations of the species occurred east of ~90° W, an area of the northern Gulf characterized as ‘high energy’ (Sturges and Leben 2000). Most NRDA and GoMMAPPS observations in this region were along the northwestern, northern, and eastern borders of the highly dynamic Loop Current, all areas especially prone to formation of eddies and detached rings (Yang et al. 2020). Black-capped petrels are therefore making extensive use of edges along a western boundary current system inside the Gulf, quite similar to its habits along the western boundary current system along the Gulf Stream in the Atlantic Ocean (e.g., Haney 1987, Simons et al. 2103, Winship et al. 2018). Conversely, only seven records of black-capped petrels were obtained from all surveys in the central and western portions of the northern Gulf of Mexico (~90 - 97° W). Although some scarcity of records here might be due to lower overall survey effort, our habitat model does not suggest this region of the Gulf contains as much suitable habitat. In fact, black-capped petrels have not been observed west of ~95.5° W despite 71 days of pelagic birding trips conducted off south Texas from 1994–2018 (https://texaspelagics.com/summary-table/), some or all of which were in deeper waters preferred by this species.

Observations of black-capped petrels in the southeastern Gulf are rare. eBird reports three boreal spring records (all in April) from inside the western Florida Straits near the Florida Keys and Dry Tortugas. A single black-capped petrel also was observed on 23 April 2011 headed northwards towards the Yucatan Channel southwest of Cozumel Island, Mexico (Haney et al. 2019). Observations of black-capped petrels in these areas would be consistent with likely migratory paths between the Gulf of Mexico and known breeding sites on Hispaniola. Nonetheless, the breeding location, breeding status, or age of black-capped petrels using the Gulf remains unknown and no data are available on connectivity between nest sites and Gulf waters. Individuals tracked from nest sites in the western Dominican Republic have not used Gulf waters (Jodice et al. 2015, Satgé et al. 2019). The closest known nesting area to the Gulf is in southern Haiti although nest sites are suspected west of that site in southeast Cuba. Our data and other records suggest black-capped petrels are present in the Gulf throughout the year, although 75% of the individuals we observed in the Gulf occurred during July – September (i.e., post-breeding phase). Our data are not, however, sufficient with respect to breeding status or age of individuals to document use of Gulf waters in relation to breeding status or age.

Our models predicted that habitat suitability for black-capped petrels increased primarily with increases in sea surface salinity (SSS), and to a lesser extent with south and eastward currents and with moderate values of sea surface height (SSH). Black-capped petrels therefore appear to be inhabiting waters that represent edges or boundaries of water masses. Spectacled petrels (*Procellaria conspicillata*) in the eastern South Atlantic also were more abundant over waters with high SSS (Camphuysen 2001). In that region, edges of Agulhas Rings (eddies specific to the conversion zone between the Atlantic and Indian oceans) are characterized by relatively high SSS and strong currents, both of which appear to concentrate prey for petrels. In the Indian Ocean, Barau’s petrel (*Pterodroma baraui*) also tend to be associated with areas characterized by levels of salinity associated with boundaries of water masses (Pinet et al. 2009). Within the Gulf of Mexico, higher levels of SSS are associated with dynamic waters associated with the Loop Current (e.g., compared to waters associated with the continental edge) particularly along the west Florida Shelf (Paluskiewicz et al. 1983) where we observed the greatest number of black-capped petrels. In the northern Gulf, the presence of squid-eating cetaceans also was associated with higher salinity waters (Davis et al. 2000). Squid also represent a primary prey item for black-capped petrels (Simons et al. 2013). The other two variables of influence identified in our Maxent models, current direction and SSH, also suggest an association with waters that likely concentrate prey. For example, Poli et al. (2017) found that SSH influenced the foraging habitats of masked boobies (*Sula dactylatra*) and posited that this feature likely serves as a surrogate for availability of forage fish. Although most of our observations occurred over the continental shelf break and slope, our models did not identify bathymetry as a strong predictor likely due to the wide range in depth that occurs along the shelf break and slope in the Gulf.

While our data are insufficient to explicitly document conservation threats to the species, we can identify threats that are likely to occur at the macro-scale (i.e., spatial and temporal overlap of a threat and species occurrence; Burger et al. 2011). The Gulf of Mexico is considered one of the marine basins with the densest activity for the extraction of oil and gas (BOEM 2012). Three potential conservation threats associated with this high level of oil and gas activity in the Gulf include collision with structures, interaction with produced waters (Middleditch 1984, Ramirez 2005), and direct impacts from oil spills. Black-capped petrels are known to collide with lighted towers and structures near breeding sites (Simons et al. 2013) and therefore the potential may exist that collision may occur with lighted structures at-sea (Montevecchi 2006). Use of waters adjacent to active oil activity by seabirds may expose individuals to produced waters which include numerous chemical constituents (Veil et al. 2004, Welch & Rychel 2004). Exposure may be direct (e.g., contact with contaminated waters; O’Hara & Morandin 2010) or indirect (e.g., ingestion of contaminated prey; Paruk et al. 2016). Petrels also may be exposed to both direct and indirect effects of oiling. Although no black-capped petrels have been recorded as mortalities from oil spills in the Gulf of Mexico, based on our data, individual petrels observed during the NRDA vessel survey were within the spatial footprint of the total slick area following the *Deepwater Horizon* blowout. Lastly, although commercial fishing activities are spatially and temporally widespread in the Gulf, the extent to which petrels overlap with these is unknown, as data with which to evaluate this threat are still relatively sparse. The species has not been historically identified in the records of pelagic observer programs although a recent study predicted that black-capped petrels may be at risk of bycatch in the pelagic longline fishery of the western north Atlantic (Simons et al. 2013, USFWS 2018, Zhou et al. 2019).

Integration of observations from NRDA and GoMMAPPS cruises and modeling in Maxent indicate that black-capped petrels are most likely to occur in the eastern region of the Gulf east of ~ 88°W (Fig. 6a) where they appear to be associated with dynamic waters of the Loop Current. The species also occurs west to ~ 95°W and south to ~ 24°N, although in a patchier distribution. Therefore, at a minimum, the northern Gulf of Mexico should be considered within the core marine range for the globally endangered black-capped petrel (Fig. 6b). Black-capped petrels occurred in all four of the marine ecoregions that comprise the Gulf of Mexico (Northern Gulf, Southern Gulf, Floridian, and Greater Antilles; Spalding et al. 2007). Nonetheless, gadfly petrels in general are highly mobile and rely on marine habitat that is highly dynamic in space and time. The definition of a marine range for such species might best be considered in terms of broad marine areas that offer habitat conditions amenable to foraging and flight. Therefore, an integration of both known occurrences and probabilistic occurrences based on habitat suitability appears to offer an approach that strikes a balance between the challenges of observing a rare and highly mobile species in remote locations and the predictive and informative nature of habitat modeling. We suggest that the range for black-capped petrels include the entire northern Gulf of Mexico with recognition that distribution may be more clumped in the eastern Gulf and patchier elsewhere. The results from models in Maxent predict there are some areas that offer suitable habitat within which we did not observe birds. Furthermore, the southern Gulf of Mexico remains under-surveyed for this species and hence represents a substantial data gap. With additional effort, areas that were modeled as suitable yet devoid of occurrences may receive additional survey attention and their status refined (e.g., southwestern Gulf).

**Fig. 6.**
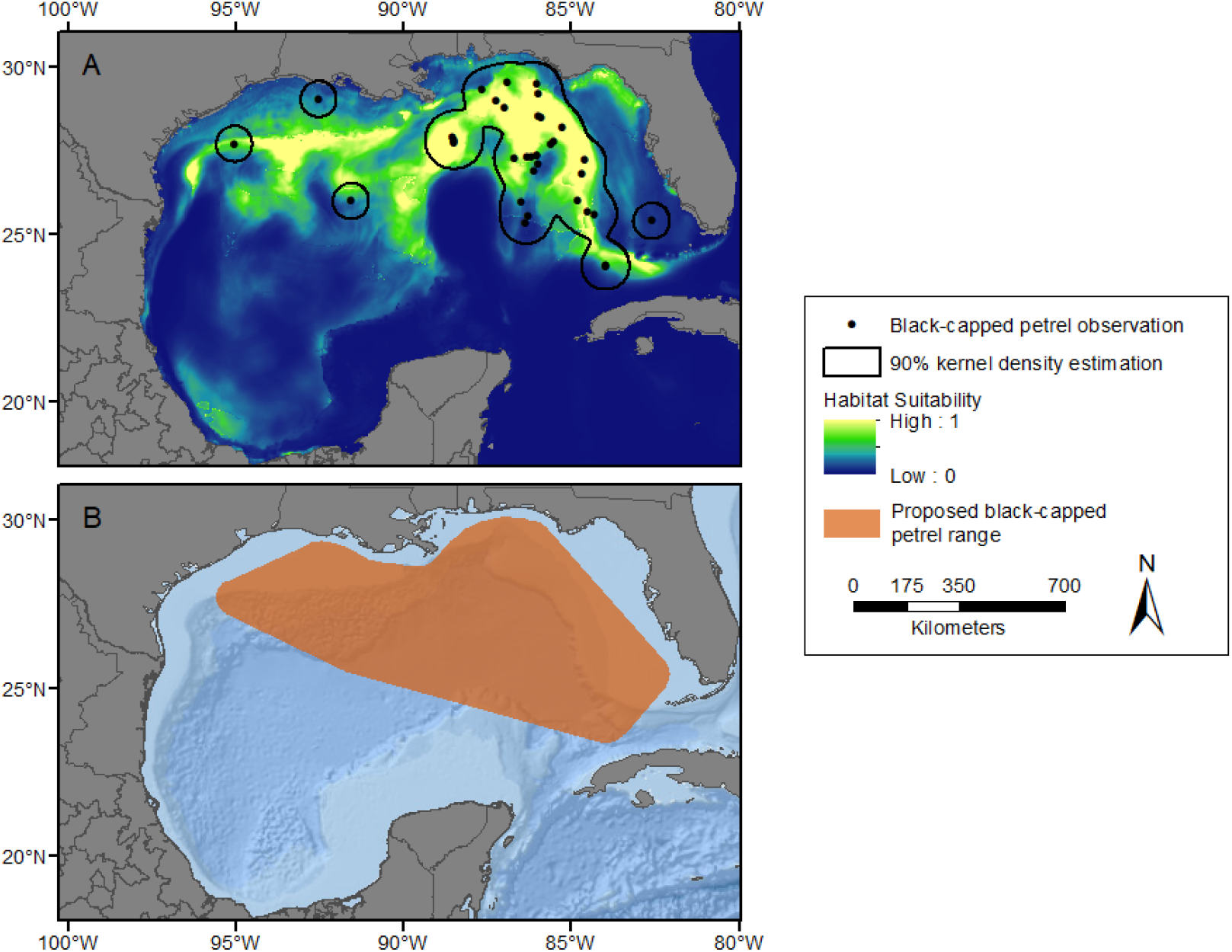
Black-capped petrel (*Pterodroma hasitata*) in the northern Gulf of Mexico. (A) 90% utilization distribution (based on observations collected from surveys conducted during the post-spill Deepwater Horizon Natural Resources Damage Assessment (NRDA) and the Gulf of Mexico Marine Assessment Program for Protected Species (GoMMAPPS)) overlaid on predicted habitat suitability, and (B) proposed range for black-capped petrels in the northern Gulf of Mexico as indicated by a minimum convex polygon encompassing the 90% utilization distribution. Credit for base map: ESRI, Garmin, GEBCO, NOAA, NGDC, and other contributors.

Efforts to better understand the marine and terrestrial range of the black-capped petrel have been increasing in recent years and that has allowed for an enhanced ability to focus research and prioritize conservation actions by stakeholders (Goetz et al. 2012). As the number of observations of black-capped petrels in the Gulf increases, a reassessment of habitat associations, model fit, and model stability would appear to be warranted and would benefit agencies responsible for regulatory actions related to anthropogenic activities (e.g., oil and gas extraction, wind-energy development, offshore fish farms), oil spill modeling and response efforts, and listing reviews. Efforts to deploy tracking devices on breeding black-capped petrels at nest sites (Jodice et al. 2015, Satgé et al. 2019) have improved our understanding of use areas, connectivity between use areas and known nesting areas, and fidelity to and residence time within use areas. To date, however, all tags deployed have been from only one nesting area in the western Dominican Republic, and none of the birds tagged were tracked to the Gulf of Mexico. Therefore, although our results suggest spatially and temporally widespread use of the Gulf, it remains unclear which of the few remaining nesting areas of black-capped petrels are directly linked to the Gulf of Mexico.

## ACKNOWLEDGEMENTS

Funding for the NRDA 2010-2011 surveys was provided through the Deepwater Horizon Natural Resources Damage Assessment administered by spill trustee Department of the Interior, USFWS. Funding for GoMMAPPS surveys was provided by BOEM to the US FWS (IAA #M17PG00011). Surveys were conducted on-board NOAA and other vessels. We are grateful to the crews and NOAA field party chiefs of the R/Vs *Gordon Gunter*, *Nancy Foster, Oregon II*, and *Pisces*, as well of the crews of UNOLS R/V *F.G. Walton Smith*, M/V *Nick Skanski*, and USGS cutter *Cypress* for their logistical support. We also thank the many seabird observers who participated in each program: Jonathan M. Andrew, Dan Bauer, Peter J. Blank, Katie Cowen, Dawn Breese, Sarah L. Flaherty, Logan Fordham, E. Wayne Irvin, Ron Goddard, Elizabeth T. Hug, Carol Keiper, David S. Lee, Matthew Love, Scott McConnell, Michelle McDowell, Nicholas Metheny, G. Scott Mills, Mark Oberle, Jim Panaccione, Stormy Paxton, Stephanie Powell, Carlos Sanchez, and Alex Wang. Kathy Hixson created and enhanced maps that appear in the figures. Raul Ramos and Paige Byerly provided helpful reviews of the manuscript prior to submission. The South Carolina Cooperative Fish and Wildlife Research Unit is jointly supported by the U.S. Geological Survey, South Carolina DNR, and Clemson University. Any use of trade, firm, or product names is for descriptive purposes only and does not imply endorsement by the U.S. Government. The findings and conclusions in this paper are those of the author(s) and do not necessarily represent the views of the U.S. Fish and Wildlife Service.

