## Supplemental Table 1 for "Expanding the marine range of the endangered black-capped petrel *Pterodroma hasitata*: Occurrence in the northern Gulf of Mexico and conservation implications"

Submission date: 17 January 2021

Table S1. Dates of all research cruises from two research programs from which observations of black-capped petrels (*Pterodroma hasitata*) were recorded in the northern Gulf of Mexico. Surveys conducted in 2010 and 2011 were a component of the post-spill *Deepwater Horizon* Natural Resources Damage Assessment (NRDA). Surveys conducted in 2017 – 2019 were a component of the Gulf of Mexico Marine Assessment Program for Protected Species (GoMMAPPS). Detections refer to an observation of ≥ 1 black-capped petrel.

| Survey program/ dates of cruise | Count of detections during cruises | Count of individuals during cruises |
|---|---|---|
| NRDA | | |
| 08 July - 12 July 2010 | 0 | 0 |
| 13 July - 18 July 2010 | 2 | 2 |
| 26 July - 02 August 2010 | 0 | 0 |
| 02 August - 07 August 2010 | 0 | 0 |
| 07 August - 21 August 2010 | 1 | 1 |
| 21 August -26 August 2010 | 0 | 0 |
| 24 August - 09 September 2010 | 1 | 1 |
| 14 September - 28 September 2010 | 0 | 0 |
| 15 September - 28 September 2010 | 0 | 0 |
| 08 October - 18 October 2010 | 0 | 0 |
| 13 October -28 October 2010 | 0 | 0 |
| 21 October - 03 November 2010 | 0 | 0 |
| 02 November - 12 November 2010 | 0 | 0 |
| 09 November - 19 November 2010 | 0 | 0 |
| 16 November - 22 November 2010 | 0 | 0 |
| 3-4, 6-12, 14 December 2010 | 0 | 0 |
| 17 February - 1 March 2011 | 1 | 1 |
| 10 March - 22 March 2011 | 0 | 0 |
| 24 March - 27 March 2011 | 0 | 0 |
| 7-15 April, 17-25 April 2011 | 0 | 0 |
| 24 April - 28 April 2011 | 0 | 0 |
| 25 April - 30 April 2011 | 0 | 0 |
| 02 May - 10 May 2011 | 2 | 4 |
| 13 May - 27 May 2011 | 0 | 0 |
| 07 June -11 June 2011 | 0 | 0 |
| 23 June - 04 July 2011 | 0 | 0 |
| 06 July - 17 July 2011 | 0 | 0 |
| GoMMAPPS | | |
| 28 April – 11 May 2017 | 0 | 0 |
| 16 May – 30 May 2017 | 1 | 1 |
| 04 June – 16 June 2017 | 0 | 0 |
| 21 July – 05 August 2017 | 3 | 3 |
| 09 August – 25 August 2017 | 5 | 5 |
| 17 September – 29 September 2017 | 0 | 0 |
| 14 January – 19 January 2018 | 0 | 0 |
| 26 January – 08 February 2018 | 0 | 0 |

| | | |
|---|---|---|
| 12 February – 26 February 2018 | 0 | 0 |
| 01 March – 16 March 2018 | 2 | 2 |
| 27 April – 11 May 2018 | 2 | 2 |
| 16 May – 25 May 2018 | 0 | 0 |
| 11 August – 28 August 2018 | 13 | 15 |
| 01 September – 18 September 2018 | 0 | 0 |
| 11 September – 30 September 2018 | 1 | 1 |
| 21 September – 06 October 2018 | 0 | 0 |
| 26 April – 10 May 2019 | 0 | 0 |
| 14 May – 24 May 2019 | 0 | 0 |
| 21 August – 06 September 2019 | 1 | 1 |
| 10 September – 25 September 2019 | 1 | 1 |
